# Unsupervised cell interaction profiling reveals major architectural differences between small intestinal and colonic epithelial crypts

**DOI:** 10.1101/2020.03.06.980243

**Authors:** Jason T. Serviss, Nathanael Andrews, Agneta B. Andersson, Ewa Dzwonkowska, Rosan Heijboer, Natalie Geyer, Marco Gerling, Martin Enge

**Affiliations:** Department of Oncology and Pathology, Karolinska Institutet, BioClinicum J6:30, Akademiska stråket 1, 171 64 Solna, Sweden; Department of Biosciences and Nutrition (BioNut), Karolinska Institutet, Hälsovägen 7, 141 83 Huddinge, Sweden; Tema Cancer, Karolinska University Hospital, 171 65 Solna, Sweden

## Abstract

Cellular identity in complex multicellular organisms is strictly maintained over the course of life. This control is achieved in part by the organ structure itself, such that neighboring cells influence each other’s identity. However, large-scale investigation of the cellular interactome has been technically challenging. Here, we develop CIM-seq, an unsupervised and high-throughput method to analyze direct physical cell-cell interactions between every cell type in a given tissue. CIM-seq is based on RNA sequencing of incompletely dissociated cells, followed by computational deconvolution of these into their constituent cell types using machine learning. We use CIM-seq to define the cell interaction landscape of the mouse small intestinal and colonic epithelium, uncovering both known and novel interactions. Specifically, we find that the general architecture of the stem cell niche is radically different between the two tissues. In small intestine, the stem-Paneth cell interaction forms an exceptionally strong and exclusive niche, in which Paneth cells provide Wnt ligands^1^. In colonic epithelium, no similar compartment exists to support stem cells, and Wnt signaling is provided by a mesenchymal cell layer^2,3^. However, colonic stem cells are supported by an adjacent, previously unrecognized goblet cell subtype expressing the wound-healing marker Plet1, which is also highly upregulated during regeneration of colon epithelium. These results identify novel cellular interactions specific for the colonic stem cell niche and suggest an additional level of structural control in the colon. CIM-seq is broadly applicable to studies that aim to simultaneously investigate the constituent cell types and the global interaction profile in a specific tissue.

## Main

Cells in higher order multicellular organisms are diverse and highly specialized. Such a level of specialization requires strict control, which is in part encoded in their spatial organization^1,4^. Single-cell RNA-seq (scRNA-seq) methods have enabled rapid advances towards determining the full complement of cellular diversity in human and mouse tissue^5^. However, no similar high throughput and unbiased method exists to chart the fine-grained structural diversity of how these cells interact, or to profile interaction-dependent changes in gene expression. To this end, we developed Cell Interaction by Multiplet sequencing (CIM-seq), a method that allows large-scale interaction profiling within a well-established scRNA-seq framework.

scRNA-seq methods generally rely on a suspension of dissociated cells where those cells which are not fully dissociated from each other (multiplets) are removed^6^. In CIM-seq, we repurpose these multiplets to determine which cells were physically attached to each other in the intact tissue. RNA-seq libraries prepared from such cell multiplets represent a mix of unknown quantities of cells that exist in the tissue, and their transcriptional profile can therefore be closely approximated *in silico* by combining scRNA-seq data from the constituent cell types. Thus, a multiplet profile can be computationally deconvoluted into fractional contributions of single cells, given a set of available singlet transcriptional profiles, and an estimation of the number of cells that constitutes the multiplet. CIM-seq determines multiplet composition in three separate stages (Fig. 1a). In the first step, we perform partial dissociation of the target tissue, followed by cell sorting of singlets and multiplets into multiwell plates and conventional Smart-Seq2 library preparation^7^. Second, we use the singlet sequence data to perform automated feature selection followed by graph-based clustering to construct a blueprint of cell types and states in the tissue. In the third step, we employ computational deconvolution to perform a maximum likelihood estimation (MLE) that determines each multiplet’s most likely cell type constituents based on the previously defined blueprint. Specifically, the MLE function uses particle swarm optimization^8^ to determine which combination of the cell types has the highest likelihood of producing the observed multiplet expression profile (see *methods* for details). Finally, by examining the cell types contained in many multiplets we can create a map of specific physical cell interaction in the intact tissue.

**Figure 1.**
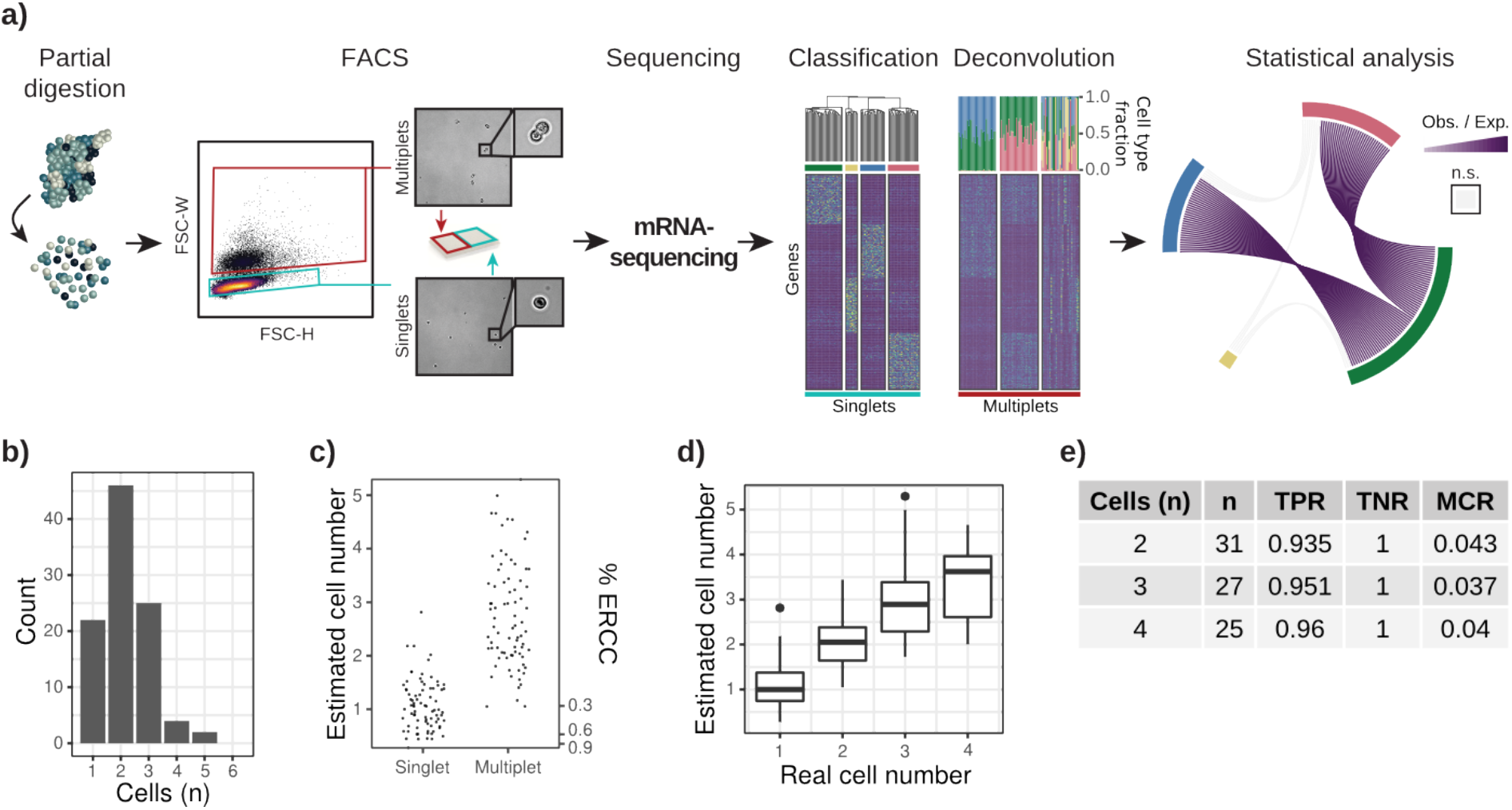
CIM-seq allows unbiased determination of physically interacting cells types. **a)** Summary of the method. Solid tissue is partially dissociated into single cells and cell multiplets, sorted separately into individual wells, and analyzed using scRNA-seq. Each multiplet is computationally deconvoluted into its most-likely single cell constituents. Statistical enrichment of co-occurring cells in a large cohort of multiplets indicates physical interaction in the tissue. **b)** Distribution of cell numbers per multiplet as determined by phase contrast microscopy. **c)** Fractional content of spike-in RNA (ERCC) can be used to estimage the number of cells in a multiplet. %ERCC (right-hand x-axis) is translated into number of cells (left-hand axis) **d)** ERCC-based cell number estimation correlates well with real cell number in multiplets with known cell composition. **e)** Error rates of multiplets with known cell composition. Mean true positive rate (TPR), mean true negative rate (TNR) and mean misclassification rate (MCR) is shown for each multiplet cell number

We first tested the propensity of cell singlets to re-associate in suspension, which would result in connections that do not reflect physical attachment in the tissue. Singlet identity remained stable over time, with singlet re-association below 0.5% after 2h (Extended Data Fig. 1a). Examining the FACS-sorted multiplets by microscopy revealed that the majority consisted of two or three physically connected cells (Fig. 1b). To measure the performance of CIM-seq in a controlled setting, we used three distinct cell lines (A375 [melanoma], HCT116 [colon cancer], HOS [osteosarcoma]) and sorted these as either singlets, or as multiplets of a known composition. The final dataset included 79 singlets and 83 multiplets (multiplets consisting of to to four cells were present in approximately equal proportions). The number of cells were estimated by comparison of cellular mRNA counts to ERCC (External RNA Control Consortium) control RNA^9^ spiked in at a known concentration (Fig. 1c,d). Cell type specific markers were observed to be exclusively co-expressed at appreciable levels in multiplets (Extended Data Fig. 1b). Dimensionality reduction (uniform manifold approximation and projection, UMAP^10^) and graph-based classification identified the individual cell types (Extended Data Fig. 1c) forming the blueprint for the deconvolution step. The deconvolution revealed a very high level of correspondence between the expected and detected connections with a <5% average error rate for each of the examined cell compositions (Fig. 1e, Extended Data Fig. 1d-1e).

To test the feasibility of performing interaction profiling in a complex tissue, we analyzed mouse small intestinal epithelial crypts of Lieberkühn, in which the organization of different cell types along the crypt-villus axis is known^11–14^. Epithelial stem cells of the intestine reside at the bottom of these crypts and their stemness is critically maintained by interaction with postmitotic, Wnt3-producing Paneth cells. Upon stem cell division, one of the daughter cells is pushed upwards towards the lumen, losing contact with the Paneth cell and thereby initiating the process of differentiation into the mature cell types of the intestine. We asked whether CIM-seq would be able to identify the Paneth-stem cell interaction in an unsupervised manner. Therefore, we sequenced 1213 single cells from mouse small intestine, and performed unsupervised classification of the scRNA-seq data, which readily identified the cell epithelial subtypes of the epithelial crypts^15^. The dimensionality reduced representation of the data showed Lgr5+ stem cells gradually transitioning into enterocytes, and correctly separated the other major cell types (goblet, Paneth, enteroendocrine, and tuft cells) of the small intestine (Figure 2a). The multiplet deconvolution relies on classes being well defined, and could potentially fail due to class ambiguity. We evaluated the quality of the classification by a) identifying differential gene expression between all of the individual classes (each class had at least 20 specific genes), and b) demonstrating that the classes are highly distinguishable by the deconvolution algorithm, by deconvoluting the singlets and recovering the correct class in 96% of the cases (Extended Data Fig. 2). In summary, these results show that the classification procedure can accurately identify specific cell types and cell states that are distinguishable by their gene expression profile.

**Figure 2.**
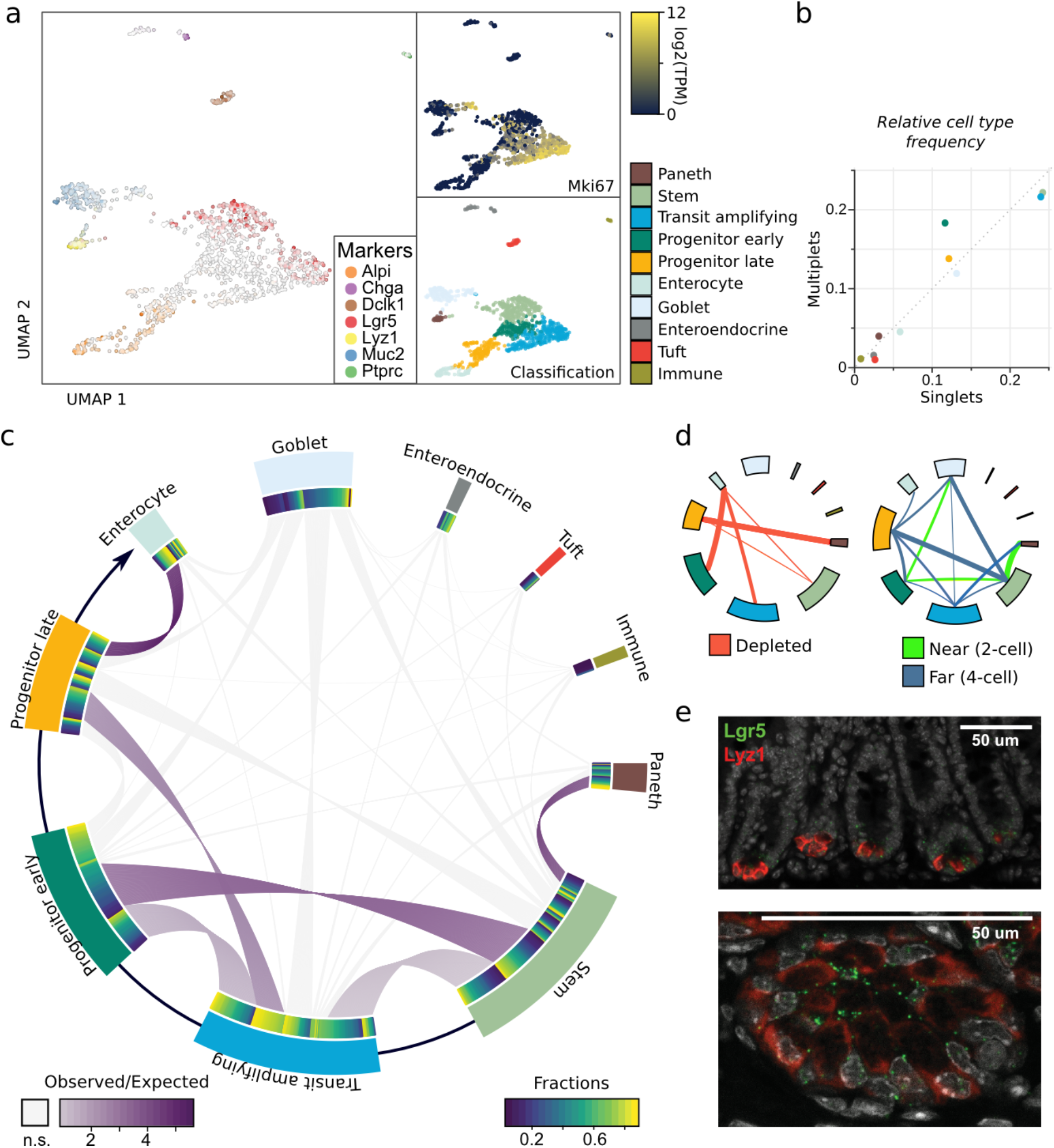
CIM-seq accurately determines the small intestinal crypt architecture. **a)** scRNA-seq analysis of mouse small intestinal epithelium. **b)** Cell type frequencies in singlets and deconvoluted multiplets. **c)** Multiplet deconvolution analysis of small intestinal crypts. Purple lines indicate that the connected cell types interact in a specific fashion (significally enriched in multiplets compared to expectation). Arrow at circumference indicate the major axis of differentiation from stem cells (bottom of crypt) to enterocytes (top of crypt). **d)** CIM-seq provides information on non-interacting cells and interaction distance. Left: Interaction depletion analysis using the same data as in c, red lines indicate significant depletion in multiplets compared to expectation, thicker lines indicate stronger depletion. Only interactions that have sufficient power to detect depletion are shown. Right: Interaction distance analysis by differential analysis of cell order. Green lines indicate relatively stronger enrichment at short distances (duplets) and blue lines at long distances (quadruplets), with line thickness indicating the strength of the relative enrichment. Note that Paneth/stem cell co-occurrence is strongly enriched at short distances, underscoring their tight bond, whereas the less ordered enterocyte/late progenitor cells are approximately equally enriched. **e)** Small intestinal stem cell niche visualized by RNA ISH. Lgr5: Stem cells, Lyz1: Paneth cells.

We subsequently performed CIM-seq deconvolution of 435 multiplets isolated from the same cell suspensions as the singlets. The distribution of cells per multiplet based on ERCC spike in ratio followed closely the distribution determined by visual inspection. Cell type frequencies inferred by multiplet deconvolution were similar to the frequencies we found in the singlet data set (Fig. 2b), indicating that there were no systematic errors in multiplet selection or deconvolution. Importantly, when the analysis was performed in conjunction with colonic tissue, no significant cross-connections were identified indicating a low frequency of false positive connections (Extended Data Fig. 2c). To find cell types with a specific preference for interaction partners, we determined the statistical enrichment of every pairwise cell type connection (see *methods* for details). Cell types that are scattered throughout the crypt, such as goblet cells, were rich in interactions but interacted with other cells at expected frequencies, and there was no depletion or enrichment of interaction with any particular cell type (Fig. 2c and 2d, left panel). Paneth cells and stem cells, in contrast, represented a highly enriched connection, with 80% of the total Paneth cell connections being to a *Lgr5*+ cell type. In line with Paneth/stem cells always being found directly adjacent to one another, this interaction was relatively more highly enriched in duplets compared to quadruplets (Fig 2d, right panel). RNA in situ hybridization (ISH) of the stem cell marker *Lgr5* and Paneth cell marker *Lyz1* confirmed the direct adjacency of the cells *in vivo* (Fig. 2e). Thus, the Paneth and stem cell connection represents a highly specific interaction in our analysis, in agreement with its role in maintaining stem cell identity by direct interaction.

The colonic epithelium has a similar crypt structure to small intestine, with Lgr5+ stem cells located at the base of the crypt and differentiating into broadly equivalent cell types (colonocytes, goblet cells, enteroendocrine, tuft). However, the architecture of the colonic crypt has previously not been characterized to the same resolution. Paneth cells, for example, are fundamental to the small intestinal stem cell niche but it is undetermined whether an equivalent cell type exists in the colon^16,17^. To directly compare the architectural properties of the two tissues, we analyzed 2467 single cells and 1703 multiplets from colon using CIM-seq. The colonic epithelium displayed a higher fraction of goblet cells, in agreement with previous observations^18^, and the goblet cells were more diverse, organizing into four different classes identified by unsupervised classification (Fig. 3a). Notably, two of these classes expressed the wound-healing marker Placenta expressed transcript 1 (*Plet1*). Even with the higher level of cell type complexity, the deconvolution-inferred cell type frequencies mirrored singlet frequencies (Fig. 3c). Connection enrichment analysis revealed a similarly strong link between terminally differentiated colonocytes and late-stage differentiating cells (Fig. 3b). Although colonocytes were strongly connected to late-stage differentiating cells, they were more highly connected to other classes in colon than enterocytes in the small intestine, potentially reflecting a less pronounced compartmentalization due to the more compact structure of the colon, which lacks the large villus structures of small intestine (Extended data Fig. 4c).

**Figure 3.**
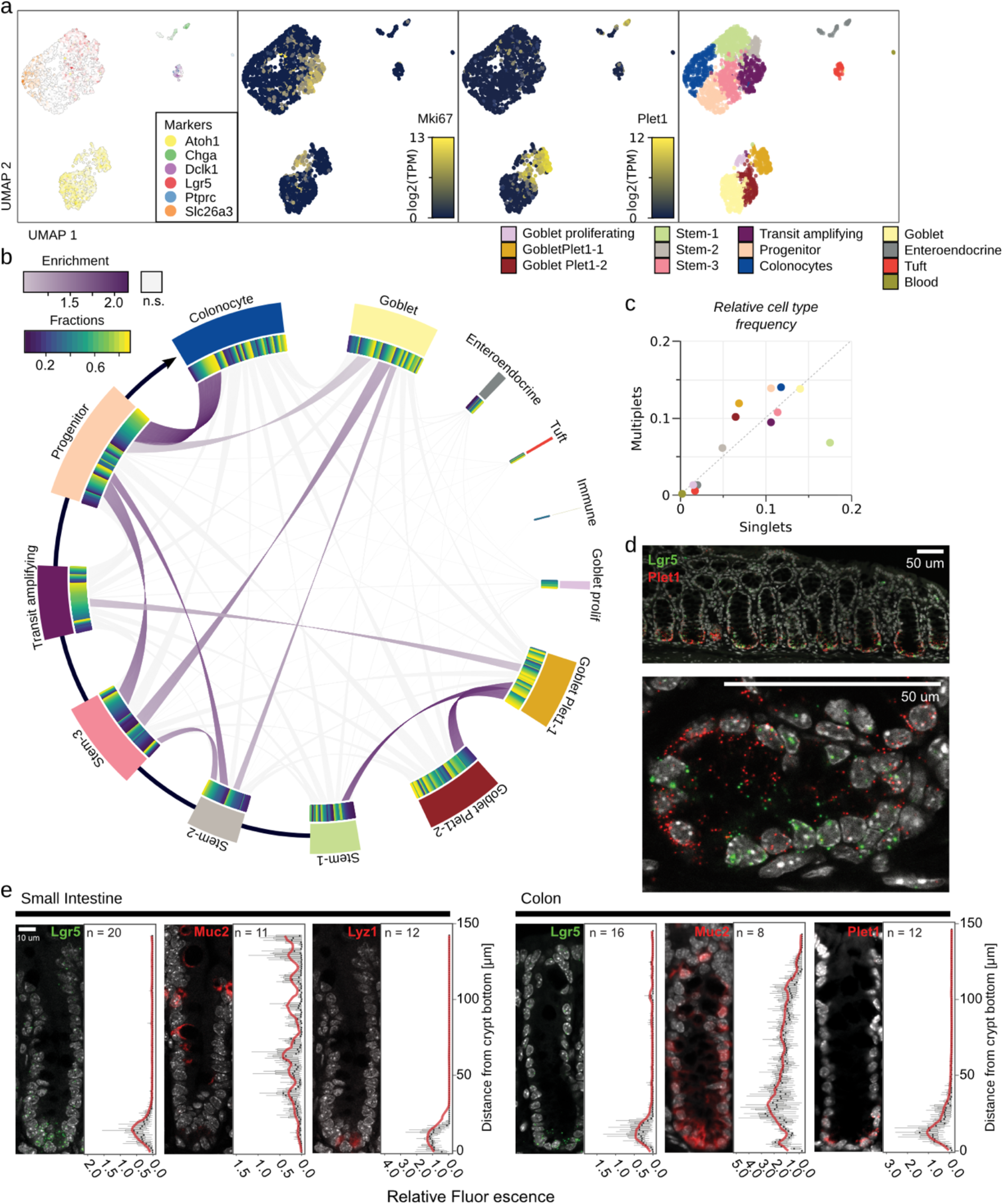
Interaction analysis of the colonic crypts reveals Plet1+ goblet cell - stem cell interaction. **a)** scRNA-seq analysis of the colonic singlets. Left: markes are Atoh1 (goblet), Chga (enteroendocrine), Dclk1 (tuft), Lgr5 (stem), Ptprc (immune), Slc26a3 (colonocyte). Middle left: Mki67 (proliferating cells). Middle right: Plet1 (Plet1+ goblet cells). Right: unsupervised classification. **b)** Multiplet deconvolution analysis of colonic epithelium. Multiplet co-occurrence is indicated by a line between classes, where significantly enriched classes are connected by purple lines, similar to Fig. 2c. Arrow connects cell types that are part of the gradual differentiation from stem cell to colonocytes, and the connected classes are sorted by *Lgr5* expression (High in Stem1, undetectable in colonocytes). **c)** Cell type frequencies in singlets and deconvoluted multiplets **d)** Detection of stem cell markers Lgr5 and Plet1 by RNA ISH of mouse colon. Top: Lgr5+ stem cells and Plet1 cells co-localize at the base of colonic crypts. Bottom: Plet1+ cells are interspersed between Lgr5+ stem cells. **e)** Distribution of stem, Paneth, goblet and Plet1+ cells in the small intestine and colon crypts, detected by RNA ISH with Lgr5, Lyz1, Muc2 and Plet1 probes, respectively. 8-16 representative crypts were analyzed by quantifying the signal intensity in vertical slices along the crypt.

The goblet cell sub-clusters displayed two distinct interaction patterns. Plet1+ goblet cells interacted strongly with the most highly Lgr5-expressing cell cluster, whereas Plet1-goblet cells preferentially interacted with more differentiated cell types (Fig. 3b). In line with this, the probability of observing *Lgr5* expression was >3-fold higher in multiplets containing goblet cells that express *Plet1* compared to *Plet1* negative counterparts (Extended Data Fig. 3b). RNA ISH verified that the Lgr5+ stem cells were located adjacent to Plet1+ cells in the intact tissue (Fig. 3d). Quantification of the RNA ISH signal confirmed localization of Plet1+ cells specifically to the crypt base, in direct proximity to Lgr5+ stem cells, mirroring the pattern of Lyz1+ cells in the small intestine (Fig. 3e). Hence, using CIM-seq we discovered a previously unrecognized cell type, characterized by the expression of *Plet1*, that directly interacts with Lgr5+ stem cells in the colon stem cell niche.

Proximity to external Wnt signaling is central to intestinal stem cell maintenance and control. In the small intestine, both Paneth cells and the basal stromal layer are thought to provide Wnt ligands to the base of the crypts, whereas for colon, the exact sources of Wnt ligands are not well defined. It has been suggested that the colon harbors a similar Wnt-expressing cell type to the Paneth cell ^16,17^, or that the colonic epithelium relies entirely on the underlying stromal layer to maintain Wnt singaling in stem cells^2,19^. When examining the expression of Wnt ligands previously reported to be important for intestinal stem cell maintenance, we found they were largely absent in the epithelium of both tissues, with the exception of *Wnt3*, which was exclusively expressed by Paneth cells of the small intestine (Fig. 4a)^19,20^. Stem cells and transit amplifying cells, in both tissues, had high expression levels of multiple Wnt target genes, indicating that Wnt signalling is active in these cells and prompting further investigation into sources of Wnt ligands^21^. RNA ISH confirmed epithelial *Wnt3* expression in small intestinal crypts, while *Wnt2b*, another Wnt ligand reported to be expressed in the intestine, was found in the stroma of both tissues, confirming previous results (Fig. 4b)^19^. Among other known factors thought to shape the intestinal stem cell niche, *Notum* had a high specificity for Paneth cells, whereas Delta-like 2 (*Dll2*) and *Dll4* were expressed by multiple cell types in both tissues (Fig. 4a)^22^. Similarly, *Reg4* and *cKit*, two other genes proposed to mark niche cells in the colonic crypt, were promiscuously expressed in all goblet cell types (Fig 4a)^16,17^. Thus, our results indicate that none of the epithelial cell types in colon provide canonical Wnt signaling, in contrast to our results in small intestine, and thereby strongly support recent results indicating that colonic Wnt ligands originate exclusively in the stroma^3^. Plet1+ goblet cells are therefore unlikely to regulate stemness via canonical Wnt activation. Nevertheless, previous studies based on the more broadly expressed markers Reg4 and cKit have provided evidence for an essential functional role of colonic niche cells in stemness maintenance^16,17^. Importantly, Plet1+ cells express both EGF and Notch ligands (Fig. 4a), which may play a role in stemness maintenance. On the other hand, given the role of Plet1 as a marker for migratory cells^23–25^, an additional type of niche control such as regulating cell motility may be important.

**Figure 4.**
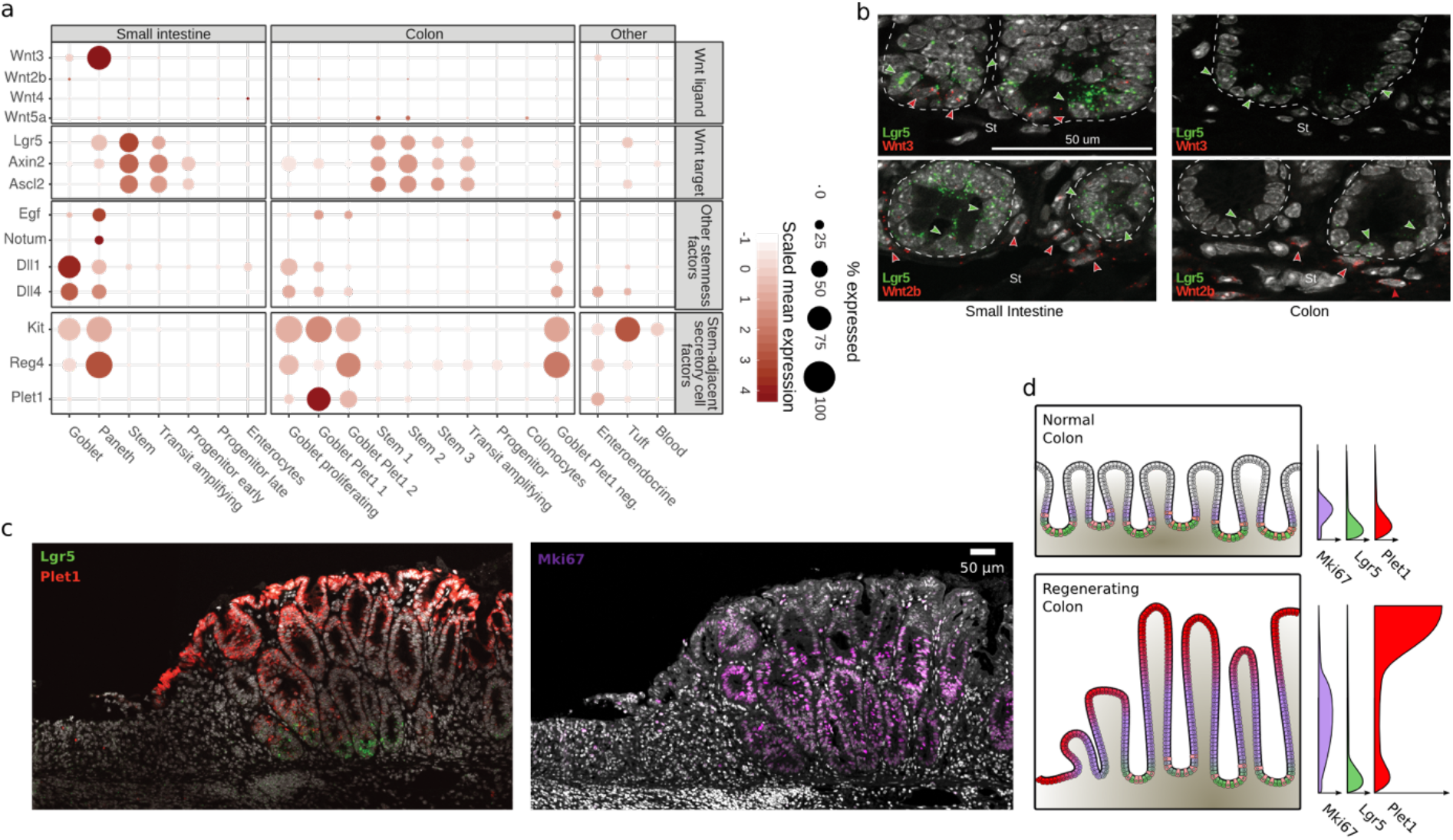
Colonic epithelium relies on external Wnt signaling, and Plet1 is highly expressed in tissue regeneration. **a)** Expression of Wnt ligands (top panel) is restricted to Paneth cells in small intestinal epithelium, whereas downstream target genes (bottom panel) are active in both tissues. **b)** Detection of Wnt3 or Wnt2a (red), and Lgr5 (green) by RNA ISH. Stroma (St). **c)** *Plet1* expression in regenerating epithelium. RNA ISH (Lgr5, Plet1) and immunofluorescence (Mki67) of mouse colon epithelium after injury induced by DSS. Region at the right is actively expanding into the damaged region (left). **d)** Schematic of Plet1-expression in colon epithelium (left panels), and expression profiles of Plet, Lgr5 and Mki67 vertically along the crypt (right panels). In normal healthy colon (top) *Plet1* and *Lgr5* are expressed at low to moderate levels in specific cell types at the base of the crypt, and proliferating cells marked by Mki67 are found in a thin layer above the base. During tissue regeneration after injury (bottom), crypts grow more elongated with a much larger layer of proliferating cells. Plet1 and Lgr5 is still expressed at the base of the crypt, but Plet1 is very strongly expressed in higher crypt regions.

*Plet1* encodes a membrane-bound glycoprotein that plays a role in cell migration after injury^23–25^ and that is functionally important for tissue repair upon damage of the intestinal epithelium. Differential expression analysis of Plet1+ goblet cells revealed genes with known roles in cell-cell adhesion (*Cgref1*, *Cd44*, *Lgals9*), as well as tissue organization and inflammation (*Ang*, *Ccl9*) to be highly expressed in Plet1+ cells (Extended Data Fig. 3a). Therefore, we next analyzed *Plet1* expression during regeneration of the colonic epithelium after chemically induced intestinal injury with dextrane sulfate sodium (DSS). We found that *Plet1* was highly upregulated in apical cells of the regenerating epithelium (Fig. 4c). By contrast, in areas of the proximal colon that remained histologically largely unaffected by DSS, Plet1 mRNA remained located exclusively to the crypt, adjacent to Lgr5 mRNA (Extended Data Fig. 4a). In regenerative areas, Lgr5+ cells were still located to the crypt bottom interspersed with Plet1+ cells, and showed increased Lgr5 mRNA levels (Extended Data Fig. 4b). Ki67 immunostaining confirmed increased levels of proliferation in regenerating crypts and indicated a shift of the proliferation zone to more luminal cells, where *Plet1* was highly upregulated (Fig. 4c, d). Together with previous data^23^, these results support a role for Plet1+ cells in intestinal damage repair and in the maintenance of intestinal tissue integrity. It is tempting to speculate that Plet1+ crypt cells could give rise to highly motile epithelial cells that repopulate damaged epithelium. Alternatively, *Plet1* is expressed ectopically by differentiated cells to aid wound repair. In either case, our results suggest that Plet1+ cells represent a highly motile cell population, which plays a role in tissue growth. Future experiments will be necessary to understand the exact nature of these cells in the stem cell niche.

Recently, several methods that allow analysis of highly multiplexed gene expression and spatial information have been developed. Multiplexed mRNA staining methods^26–28^ generally provide very high spatial accuracy, allowing for cellular or even sub-cellular localization of transcripts, but can only interrogate a number of predefined genes. Importantly, the constraint that the set of genes to measure is determined beforehand limits their use to testing predetermined hypotheses rather than generating novel ones, although higher multiplexing allows a large set of hypotheses to be tested in parallel. Array based methods are not limited to a predefined set of genes, but typically have a lower spatial accuracy due to the size of the barcoded features and diffusion rate of mRNA molecules precluding true single-cell accuracy^29,30^. Importantly, these methods are based on proximity rather than physical attachment which might make them less likely to identify permanent cell interactions. A method based on manually separating interacting cells and performing scRNA-seq on each interacting partner has also been proposed^31^. While providing excellent data on direct interaction, the reliance on specialized equipment and the high labor intensity of this method makes it impractical for general use.

CIM-seq solves many of the problems of previous methods. All transcripts are measured, similarly to index-based methods, but with the advantage that multiplets are investigated with no loss of sensitivity compared to conventional scRNA-seq. Since CIM-seq relies on actual physical attachement of cells in intact tissue, it has single-cell spatial accuracy. Also, it is based on a widely used and easy to automate scRNA-seq protocol, making it easy for labs to adopt at a large scale. Its major limitation is that it only allows us to obtain information on direct interactions and cannot detect higher-order structures, which may be important for organ development by, for example, creating a concentration gradient of signaling molecules in the tissue.

Detection of cell type interaction relies on statistical enrichment analysis to identify the type of preferential interactions between two cell types suggestive of functional co-dependence. Highly specific and exclusive interactions between transcriptionally different cells, such as the Paneth/stem cell interaction in small intestine are evident even without formal statistical enrichment analysis. However, most interactions are specific but non-exclusive (e.g. the interaction between Plet1+ goblet cells and Lgr5+ stem cells in colon) and for these, statistical enrichment analysis is necessary to account for the vast differences in cell type frequency. This also means that cell types that interact broadly in a tissue (e.g. goblet cells in the small intestine) are easily identifiable as having many non-significant interaction partners where the number of connections is well correlated with cell singlet frequency.

CIM-seq is a general method, broadly applicable to any solid tissue, and we expect it to be useful in a wide variety of scientific questions. This includes organ development and developmental diseases, decrease of fitness in aging which has been partly attributed to loss of organ integrity, and tumor-stromal interactions in malignancies. Thus, we anticipate that CIM-seq will be a useful tool to generate novel hypotheses about how specific cell interactions influence cell function and identity.

## Methods

### Cell lines

All cell lines were purchased from ATCC, verified and mycoplasma tested before use. A375 cells were cultured in Dulbecco’ s Modified Eagle’ s Medium, HCT116 cells in McCoy’ s 5a Medium Modified, and HOS cells in Eagle’ s Minimum Essential Medium (Thermofisher). All culture media were supplemented with 10% fetal bovine serum and 100U/ml Penicillin-Streptomycin (Thermofisher). Cell lines were cultured at 37°C with 5% CO2.

### Mice

C57BL/6J mice eight months of age or older were used for isolation of small intestinal and colonic epithelial cells. Breeders were bought from Scanbur, Sweden. Mice were housed in specific-pathogen-free conditions with water and standard chow diet provided ad libitum. Night-day cycles were 12 hours. All experiments and breedings were approved by the local animal ethics committee (The Board of Agriculture, Sweden).

### Intestinal injury model

DSS (approximate molecular weight 40kDa, DB001-38; Tdb Consultancy, Sweden) was initiated at a concentration of 3.5% w/v and administered in drinking water day 1 (d1) to d5. Animals were sacrificed on d8, colons were harvested, fixated in 10% w/v buffered formalin (158127; Sigma-Aldrich, USA) at room temperature overnight and subjected to paraffin embedding.

### Crypt isolation for CIM-Seq

Small intestine and colon were removed from C57BL/J wild-type mice and kept on ice in phosphate buffered saline (PBS). Lumina of colon and small intestine were washed three times with PBS and adipose tissue connected to the exterior of the small intestine or colon was removed. Colon and small intestine were opened longitudinally, and any remaining mucous on covering the epithelial layer was gently rubbed off. Tissues were washed once in PBS before being cut into 0.5 - 1 mm long fragments. Colon fragments were immersed in 10mM EDTA-PBS and incubated for 105 minutes on ice, with shaking every 15 minutes. Small intestinal fragments were similarly immersed in 10mM EDTA-PBS on ice and shaken. After 15 minutes, small intestinal fragments were allowed to settle at the bottom and the supernatant was discarded and replaced with new 10mM EDTA-PBS and shaken vigorously. This procedure was repeated 1-2 times, with supernatant fractions investigated through light microscopy before discarding, in order to enrich for crypts. Small intestinal fragments were then incubated on ice for 45-60 minutes for a total of 105 minutes. Following EDTA-PBS treatment fractions were triturated 10-15 times. Colon fractions were strained through a 100 um filter while small intestinal fractions were strained through a 70 um filter. Fractions were centrifuged at 300g for 5 minutes and dissociated using TrypLE Express (Invitrogen) at 37°C for 15-20 minutes. Using a non-specific protease such as Trypsin is recommended over a specific one (eg. collagenase) to avoid biasing multiplet composition. Enzymatic dissociation was supervised using light microscopy during regular intervals to obtain an appropriate amount of single cells and multiplets. Dead cells were removed using Dead Cell Removal Kit (Miltenyi Biotec) according to the manufacturer’ s protocol before staining for FACS sorting was performed.

### Cell sorting

Cell lines were stained with Fixable Viability Dye eFluor™ 450 (eBioscience) and live cells were sorted into 96-well plates. For synthetic multiplet generation a given number of cells from each cell line was sorted into each well for a total of 2, 3 or 4 cells. A total of 9 combinations was produced (Extended Data Fig. 4a).

Cell suspensions obtained from SI or colonic crypts were stained with FITC anti-CD326 and 7AAD (Sony Biotechnology) in order to monitor number of epithelial cells and select live cells. Additionally, Paneth cells were enriched using Brilliant Violent anti-CD24. Single cells were distinguished from multiplets using FSC-H and FSC-W gating and the gating scheme was visually confirmed via phase contrast microscopy (Fig. 1a). Cells were sorted into 384-or 96-well plates containing hypotonic lysis buffer using a SH800S FACS sorter (Sony). Plates were sealed immediately after sorting with microseal F foil (Biorad) and centrifuged at 4000g at 4°C for 5 min before being frozen on dry ice and stored at −80°C.

### scRNA-seq

Single-cell and multiplet RNA-Seq libraries were generated as described8. Briefly, single-cells collected in 384-well plates were lysed, followed by reverse transcription with template-switch using an LNA-modified template switch oligo to generate cDNA. After 21 cycles of pre-amplification and purification, DNA from representative wells was analysed on 4150 tapestation (Agilent) in order to assess cDNA quality. Tagmentation was performed using Tn5 synthesized from a psfTn5 plasmid (Addgene plasmid #79107) assembled with standard Illumina Tn5 adapters. Barcoded libraries were pooled and subjected to 75 bp single read sequencing on the Illumina NextSeq 550 instrument.

Sequencing reads were trimmed, adapter sequences removed and the reads aligned to the hg38 (human) or mm10 (mouse) reference assembly using STAR^32^ with default parameters. Duplicate reads were removed using picard^33^. Transcript counts were obtained using HTSeq^34^ and hg38 or mm10 UCSC transcript annotations. Transcript counts were normalized into log transformed counts per million. Single cell profiles with the following features were deemed to be of poor quality and removed: 1) cells with less than a specified total number of valid counts in exonic regions (1.8×10^4^ and 4×10^4^ for the mouse gut and sorted multiplets dataset, respectively), 2) cells with low actin TPM, and 3) cells with high fractions of ERCC reads. To determine a cut-off for actin TPM, we used the normal distribution with empirical mean and standard deviation from log2 transformed actin TPM, which is normally distributed in successful experiments. The cut-off was set to the 0.01 quantile (eg. the lower 0.01 % of the bell curve). A similar strategy was used for fractions of ERCC reads, samples with a log2 transformed fraction of ERCC reads above the 0.99 % quantile were rejected. In total 3680 singlets (out of 4125 analyzed wells, 89% success rate) and 2138 multiplets (out of 2452, 87% success rate) were retained for further analysis in the mouse gut dataset. In total 79 singlets (out of 96 analyzed wells, 82% success rate) and 83 multiplets (out of 96, 86% success rate) were retained for further analysis in the sorted multiplets dataset.

### CIM-seq

CIM-seq is implemented in the R statistical programming language^35^ and is freely available at (https://github.com/EngeLab/CIMseq) under a GPL-3 license. The CIM-seq method consists of three distinct stages; a preparatory stage, including feature selection, dimensionality reduction, and classification, followed by a deconvolution stage, and an interaction enrichment testing stage.

#### Preparatory stage

CIM-seq permits the elements of the preparatory stage to be carried out using appropriate methods selected by the end user. For the analysis of the data included here we choose to utilize the Seurat^36^ package version 2.3.4 for all three preparatory stage elements; i.e. feature selection, dimensionality reduction, and classification. The standard Seurat workflow was followed but briefly outlined here. Dimensionality reduction for visualization was performed using the UMAP algorithm. Unsupervised classification of cell types and cell states was performed using a graph-based method. Briefly, data was scaled and centered using the *ScaleData* function after which feature selection was performed using *FindVariableGenes* function, selecting only those genes with a mean expression of 0.5 or 1 and dispersion > 1 and otherwise using default parameters. Principal component analysis with the resulting features was performed followed by jack straw testing to determine significant (p < 0.001) principal components. Significant principal components were subsequently used as input to the graph-based classification algorithm via the *FindClusters* function. Specifically, the louvain community detection algorithm was utilized to detect cell types with 100 random starts. Classification was often performed in an iterative fashion with various settings of the resolution parameter with final classifications judged to be sufficient based on the partitioning of known cell types and marker genes. Feature selection for downstream deconvolution was carried out via the *FindAllMarkers* function using the AUC test and setting the minimum difference in the fraction of detection parameter to 0.2 or 0.4 and log fold change threshold to log(2).

#### Deconvolution stage

In the deconvolution stage the goal is to determine the fractional contribution of the various cell types, discovered in the preparatory stage, to each individual multiplet and, in this way, determine their cellular composition. The deconvolution takes advantage of particle swarm optimization (PSO) where [0, 1] constrained candidate solutions (swarm particles), consisting of a vector of fractions with one value per cell type, are optimized with respect to a cost over a number of iterations. PSO makes few assumptions about the problem being optimized, and does not require a differentiable optimization problem, while still being able to search a large space for candidate solutions^37^.

The CIM-seq cost function is based on the probability, p(m | s,c), of observing the gene expression profile of a multiplet (m), given a candidate solution (s), and the singlet gene expression profiles from each class (c). The probability is determined empirically by creating a number of *in silico* multiplet profiles derived from gene expression values of one randomly selected singlet from each class (cv, synthetic multiplets), multiplied by the candidate solution vector (s), and summed over all classes. Each of these gene expression point values are treated as individual poisson processes with λ = round(s * cv) with the joint point p-value (probability mass function, pmf) for a gene j in m (m_j_), over n randomly generated multiplets given by:

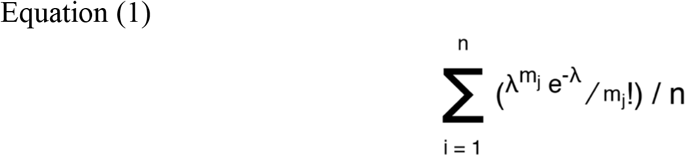

The cost is defined as the sum of the −log10 probabilities, p(m | s,c), given by Equation (1). The final result of the deconvolution procedure gives one solution, a vector of fractions, per multiplet and a corresponding cost.

Converting the solution vector of fractions into connections was achieved by first normalizing the elements of the solution vector corresponding to each cell type by the ERCC-estimated median relative RNA contribution for each cell type. Fractions are then multiplied by the ERCC-estimated cell number for each multiplet to take into account the number of estimated cells in the multiplet and then subsequently rounded to the nearest integer. Due to the inherent noise in the ERCC data, outlier multiplets (< 5%) will appear to contain an exceptionally large number of cells. In order to limit false positive connections in the deconvolution results, the estimated cell number in the multiplets was limited to 4, based on microscope observations (Fig. 1b). The resulting matrix from this procedure was then binarily transformed by converting all integers greater than 0 to 1 which indicates a connection between the corresponding cell types.

CIM-seq uses a modified SPSO2007 reference implementation of the PSO algorithm with two additions designed to improve performance when used in conjunction with CIM-seq. These modifications include 1) increased user control over early stopping criteria allowing for early termination of the optimization and 2) acceptance of user supplied swarm particle starting positions. For 1) we allowed early termination in the case that the cost did not improve by 1 (costs tend to be on the scale of thousands) in 5 iterations for all deconvolutions. For 2), we supplied precalculated swarm positions for all possible cell class combinations of one and two and added random normally distributed noise with a mean of 0 and standard deviation of 1 / the number of cell classes. In each of the deconvolutions we allow a maximum iteration of 100 with a minimum of 400 swarm particles and 2000 (Mouse gut dataset) or 400 (sorted cell lines dataset) synthetic multiplets are provided to the cost function.

#### Interaction enrichment testing stage

We assume that if connections are randomly distributed between cell types that the number of connections between two individual cell types would follow the relative abundance of those cell types. Therefore to calculate the expected number of connections between any two cell types, we first estimate the relative abundance of each cell type based on the data from the deconvoluted multiplets. Subsequently, for each cell type, we calculate the expected number of connections between that cell type and every other cell type by multiplying the relative abundance of each of the other cell types by the total number of detected edges from the cell type under consideration. The enrichment score for a specific interaction is then calculated as the quotient of the number of observed edges and the number of expected edges.

Hypothesis testing to evaluate the probability of observing a greater number of interactions than the observed number, P[I > i], is evaluated using the hypergeometric distribution. The hypergeometric distribution is commonly used to identify under-or over-represented subpopulations within a sample and is suitable to the given circumstances due to the fact that it describes the probability of *n* successes in *y* draws without replacement. Testing is performed for each interaction with test parameters set as follows where cell type one and two are defined as Ct1 and Ct2, respectively:

Population size: the total number of detected cell types in the all multiplets
Number of success states in the population: the abundance of Ct2 detected in the multiplets
Number of draws: total number of detected Ct1 interactions
Number of observed successes: number of Ct1-Ct2 interactions

The lower tail probability, P[I > i], is calculated for all interactions and probabilities are subsequently FDR corrected. H_0_ for testing is that the true number of interactions are less than or equal to the observed value. H_0_ was rejected when FDR corrected p-values were < alpha, with alpha = 1e-3 for all analyses in the study. Connections resulting from the deconvolution of the mouse gut dataset were furthermore required to have a weight, i.e. the number of multiplets with said connection, >10 to be reported as enriched.

### RNA in situ hybridization (RNA ISH) and immunofluorescence

The RNAscope Multiplex Fluorescent Reagent Kit v2 (cat nr 323100, Bio-Techne Ltd, UK) was used for RNA ISH according to the manufacturer’s instructions. In case of very high expression of a target (Lyz1, Muc2), the probe was diluted 1:10 in probe diluent (cat nr 300041) prior to hybridization. Paraffin-embedded samples fixed with either 4% paraformaldehyde or 10% buffered formalin, cut in 4 μm sections were used for all stainings.

Probes were as follows:

Mm-Lgr5-C2 (312171-C2)
Mm-Lgr5 (312171, C1 and C2)
Mm-Plet1 (557941)
Mm-Wnt3a-C2 (405041-C2)
Mm-Muc2-C2 (315451-C2)
Mm-Wnt2b-C2 (405031-C2)
Mm-Lyz1-C2 (415131-C2)
Mm-Alpi (436781)
Mm-Slc26a3 (593261)

For IF, paraffin-embedded sections were deparaffinized, rehydrated, and antigen retrieval was done with target retrieval solution (Target Retrieval Solution, Dako, Denmark, S1699) according to the manufacturer’s instructions.

Stainings were performed with the following antibodies in TBS-T (Tris Buffered Saline, 0.05% Tween 20, 1% BSA, 10% normal goat serum, 0.05% Triton X-100):

Rabbit-Anti-Phospho-Histone H3 (Ser10), Millipore #06-570, concentration 1:300 Rabbit-mAb-Ki67(D3B5), Cell Signaling, #12202, concentration 1:400

Antibody incubation was allowed overnight at 4°C, before washing and incubation with the secondary antibody (Invitrogen, Alexa Fluor 647 goat-anti-rabbit, A21245, concentration 1:400) for 1 h at room temperature.

A Zeiss LSM710 confocal microscope in single photon mode was used for imaging of mRNA ISH and IF. Nuclear counterstain was done with DAPI.

### Image quantification

Images of small intestine and colon epithelium were taken at 20x magnification. Subsequent analysis was performed using ImageJ. Crypts were manually selected and cropped. Cropped images were split into separate channels displaying signal for either DAPI or the fluorescent probe of interest. A signal threshold was set for each channel, removing noise, and channels were converted to binary where each pixel is determined as either positive or negative for fluorescent signal. Binary images were split vertically into 100 equally large segments and positive area was measured for each segment. Fluorescent signal of the probe of interest was normalized by DAPI signal from corresponding segment.

