## Supplementary figures 1-4 for "Unsupervised cell interaction profiling reveals major architectural differences between small intestinal and colonic epithelial crypts"

### Extended data

a)

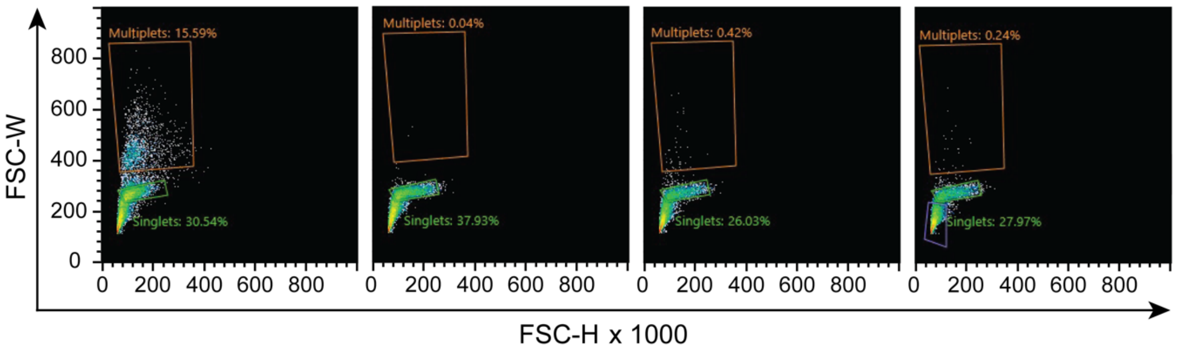


b)

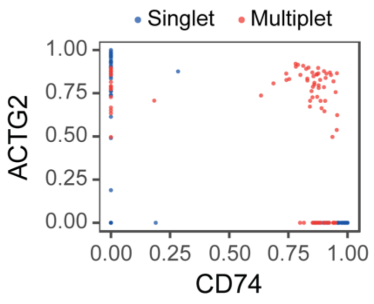


c)

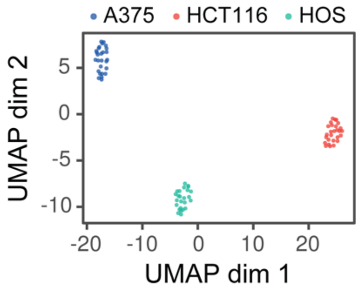


d)

| Cell type | Expected | Detected |
|---|---|---|
| A375 | 65 | 63 |
| HCT116 | 54 | 50 |
| HOS | 65 | 61 |

e)

| Cells in multiplet | n | TPR | TNR | MCR |
|---|---|---|---|---|
| A375, A375, HCT116, HCT116 | 8 | 1.00 | 1 | 0.00 |
| A375, A375, HCT116, HOS | 8 | 0.88 | NA | 0.12 |
| A375, HCT116 | 10 | 0.90 | 1 | 0.07 |
| A375, HCT116, HOS | 10 | 0.97 | NA | 0.03 |
| A375, HOS | 11 | 0.91 | 1 | 0.06 |
| A375, HOS, HOS | 9 | 1.00 | 1 | 0.00 |
| A375, HOS, HOS, HOS | 9 | 1.00 | 1 | 0.00 |
| HCT116, HCT116, HOS | 8 | 0.88 | 1 | 0.08 |
| HCT116, HOS | 10 | 1.00 | 1 | 0.00 |

**Extended Data Figure 1. a)** Flow cytometry analysis of re-association rate. HCT116 singlets and multiplets were sorted separately and singlets were re-analyzed after 0, 30, and 120 minutes. **b)** Analysis of A375 (CD74) and HOS (ACTG2) specific marker expression in singlets and multiplets shows co-expression is only observed in multiplets. **c)** Dimensionality reduction (UMAP) and unsupervised classification of the sorted cell line singlets. **d)** Numbers of expected (sorted) and detected (deconvoluted) cell types in cell line-based multiplets of a known composition. **e)** Cell line-based multiplets of a known composition results showing the number of samples (n), true positive rate (TPR), true negative rate (TNR), and misclassification rate (MCR) for each multiplet composition.

**a)**

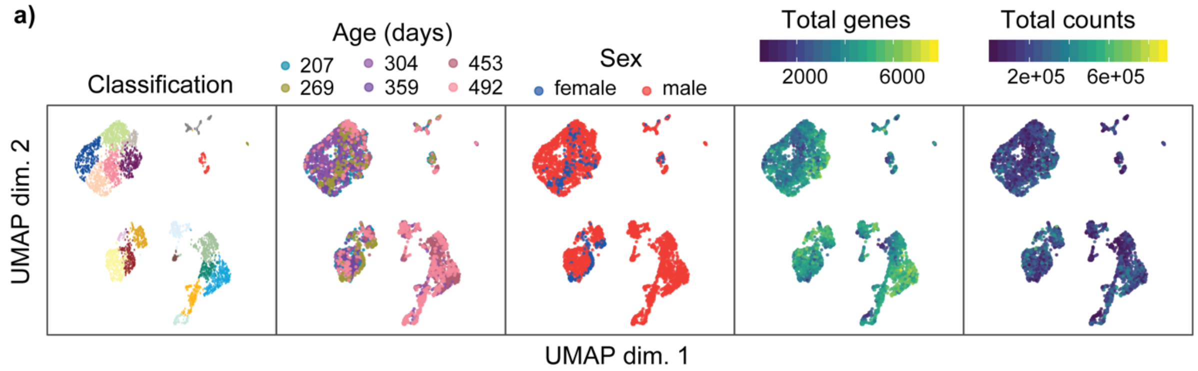


**b)**

| | Small Intestine | | | | | | Misc. | | |
|---|---|---|---|---|---|---|---|---|---|
| Class | Stem | Goblet | Paneth | Progenitor early | Progenitor late | Transit amplifying | Tufft | Blood | Enteroendocrine |
| Precision | 0.982 | 1.000 | 1.000 | 1.000 | 1.000 | 1.000 | 0.952 | 1.000 | 1.000 |

| | Colon | | | | | | | | | |
|---|---|---|---|---|---|---|---|---|---|---|
| Class | Transit amplifying | Stem 2 | Stem 3 | Colonocytes | Goblet Plet1 2 | Stem 1 | Progenitor | Goblet Plet1 1 | Goblet Plet1 neg. | Goblet proliferating |
| Precision | 0.905 | 0.931 | 0.954 | 0.966 | 0.967 | 0.967 | 0.972 | 0.983 | 0.995 | 1.000 |

**c)**

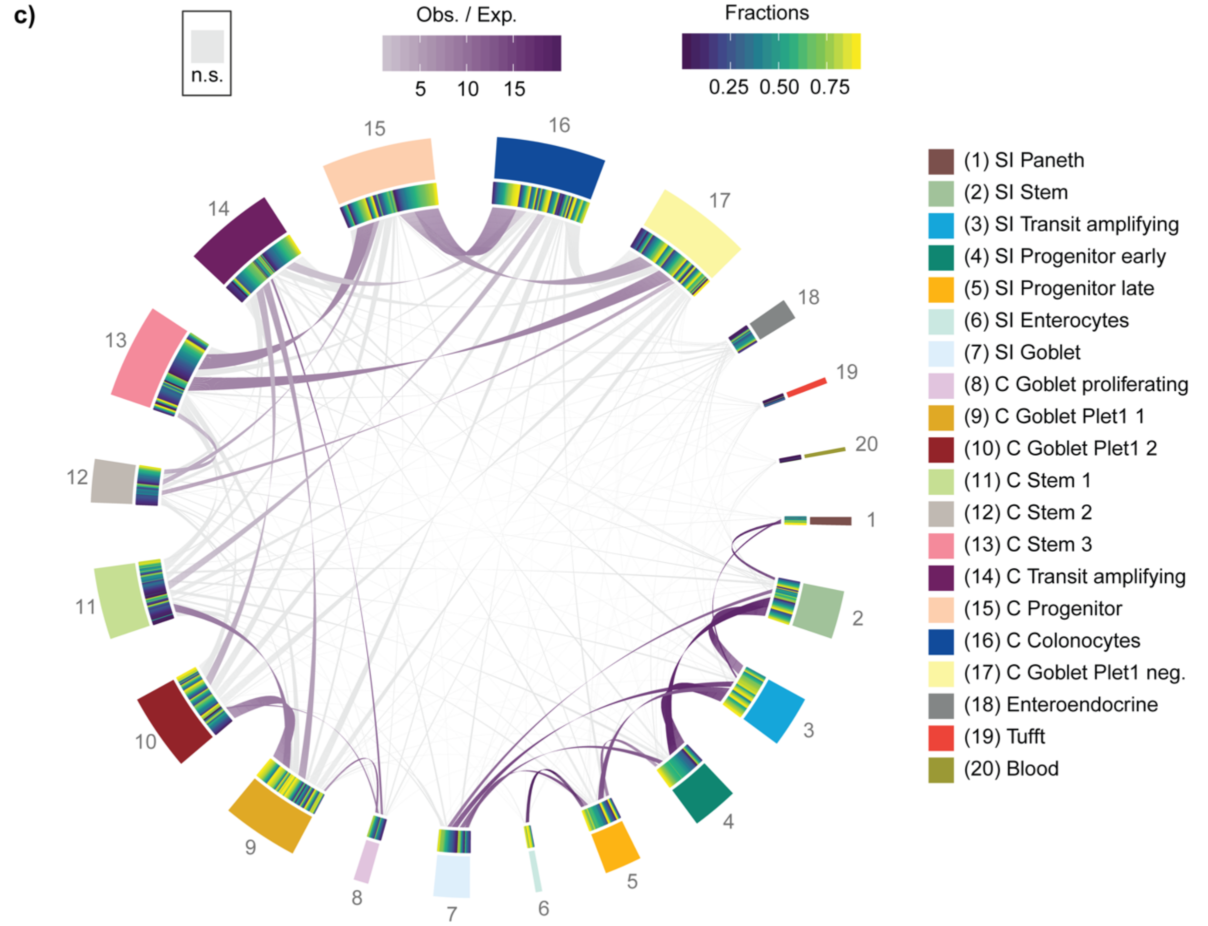


**Extended Data Figure 2. a)** Covariate analysis in mouse gut dataset showing a lack of correlation between covariates and classification. **b)** Deconvolution of mouse gut singlets shows a high precision for all classes indicating the validity of the classification and sufficiency of the provided features to allow discrimination between the different cell types. **c)** Deconvolution of the entire mouse gut dataset indicates a lack of enriched cross-tissue connections and thus implies a low false positive connection rate. Note that due to tissue-specific interactions, some previously non-significant connections (eg. goblet cells in small intestine) are now called as significant.

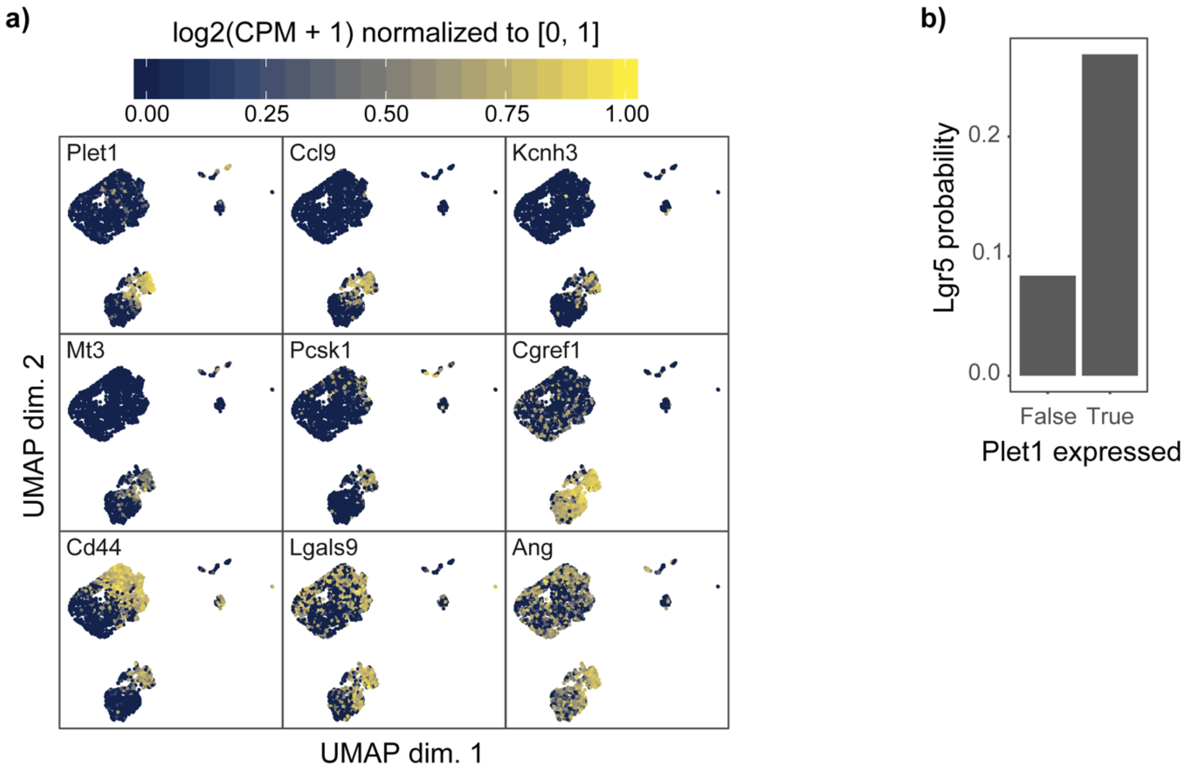


**Extended Data Figure 3. a)** *Plet1* differential expression genes, log2(TPM + 1) normalized to [0, 1], overlaid on the colonic epithelium UMAP. **b)** The probability of observing *Lgr5* expression in Muc2+ goblet cells dependent on *Plet1* expression.

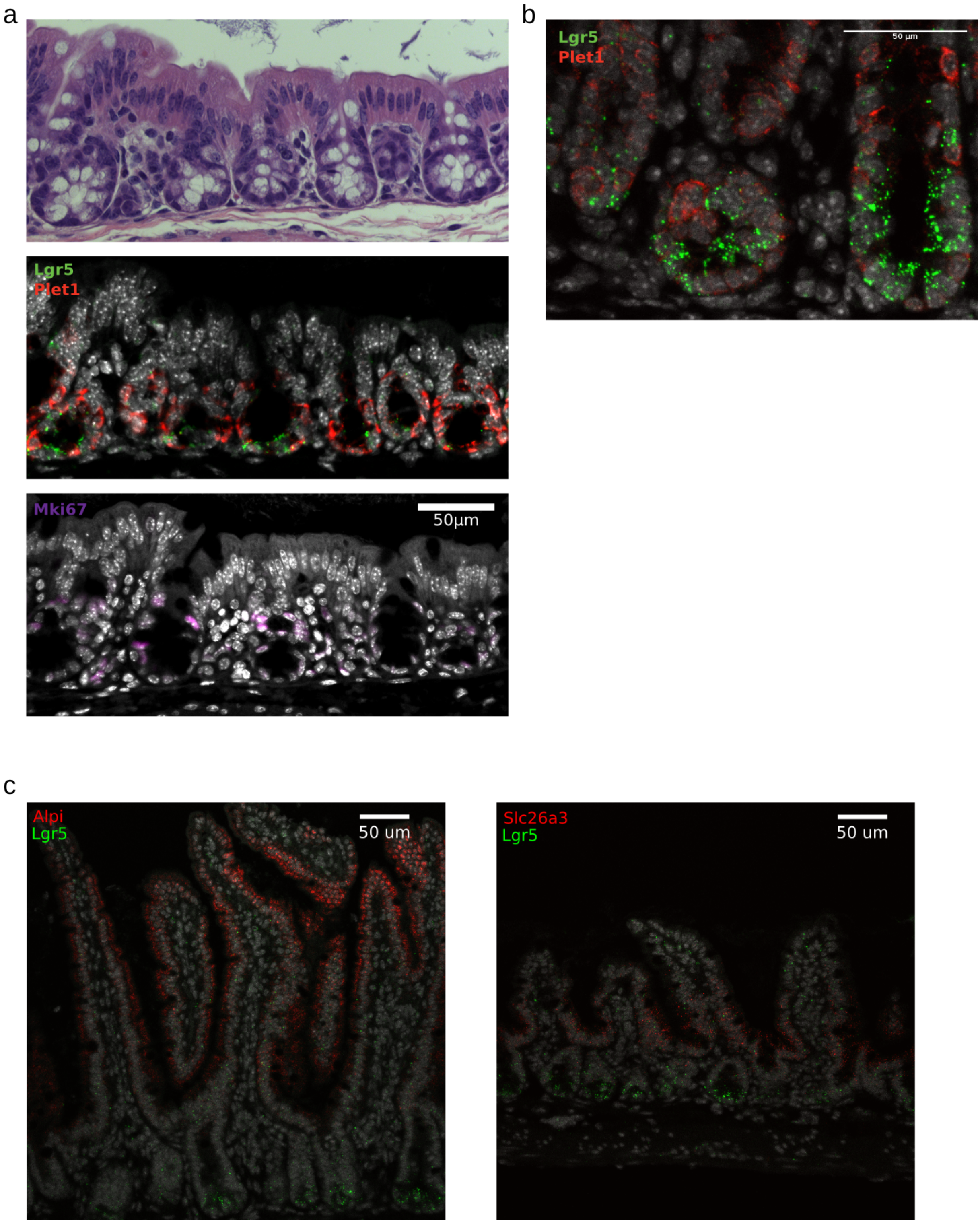


**Extended Data Fig. 4. a)** Consecutive slices from histologically unaffected regions of colon from a mouse treated with DSS. H&E (top), RNA ISH of *Lgr5*, *Plet1* (middle), and immunohistochemistry of Mki67 (bottom). **b)** mRNA abundance of *Lgr5* and *Plet1* at the bottom of crypts in regenerating colonic epithelium shows a similar pattern to that seen in normal healthy colon crypts. **c)** RNA ISH of stem cell distant cell types. Left: *Alpi* and *Lgr5* RNA ISH in small intestine. Large Alpi+ structures are villi. Right: *Slc26a3* and *Lgr5* RNA ISH in colon.
